# Facilitating Mindfulness Training with Ultrasonic Neuromodulation

**DOI:** 10.64898/2026.03.10.710890

**Authors:** Brian Lord, Erica N. Lord, Jessica Schachtner, Laura Beaman, Shinzen Young, John J. B. Allen, Joseph L. Sanguinetti

## Abstract

Systematic focus training is increasingly recognized for its therapeutic benefits, with equanimity, the ability to maintain an open and accepting attitude towards all experience, identified as a critical mechanism for improving well-being. Physiologically, experienced meditators demonstrate reduced activity in the posterior cingulate cortex (PCC), a hub of the default mode network (DMN), and increased segregation between the DMN and the central executive network (CEN). This study investigated whether non-invasive neuromodulation could facilitate these neural and behavioral shifts in novice practitioners. We conducted a single-blind, randomized controlled trial with 24 meditation-naïve participants who engaged in a two-week “Body Focus” mindfulness training program. Participants were randomized to receive either active (n=16) or sham (n=8) suppressive transcranial focused ultrasound (tFUS) targeting the PCC during four in-person meditation sessions. Resting-state fMRI analysis revealed a robust Condition × Session interaction in functional connectivity. While the sham group showed a trend toward increased coupling, the active tFUS group demonstrated significant decoupling (increased segregation) between the DMN and CEN, a pattern characteristic of advanced meditators. Subnetwork analysis indicated these effects were driven primarily by the decoupling of the core self-referential system in DMN (DMN_A_) from the external-oriented control system of the CEN (CEN_B_). Behaviorally, greater reductions in DMN-CEN connectivity within the active group predicted larger increases in self-reported acceptance and longer duration of voluntary meditation practice. These findings suggest that tFUS targeting the PCC can acutely redirect neuroplastic trajectories during early mindfulness training, potentially accelerating the acquisition of equanimity and distinct network configurations associated with effortless awareness.

**Significance Statement:** This study demonstrates that transcranial focused ultrasound (tFUS) targeting the posterior cingulate cortex (PCC) can synergistically enhance mindfulness training in novices. By inducing a robust decoupling between the default mode and central executive networks – a neural signature typically acquired only after hundreds of hours of practice – this intervention effectively redirected the neural trajectory of novice practitioners toward that of experienced meditators in just two weeks. These findings suggest that targeted neuromodulation can bypass early obstacles in meditation practice, offering a promising “precision wellness” avenue for accelerating the acquisition of equanimity and its associated well-being benefits.

## Introduction

Systematic focus training (e.g., mindfulness, meditation, and contemplative practice) has surged in popularity during the 21st century, finding application in various contexts, notably in therapeutic techniques including mindfulness-based stress reduction (MBSR) and mindfulness-based cognitive therapy (MBCT) (Alsubaie et al., 2017; Dimidjian & Segal, 2015; Sharpio & Weisbaum, 2020; Van Dam et al., 2018). These techniques have gained prevalence as an evidence-based therapeutic tool for improving physical and mental health outcomes for numerous conditions, including depression, anxiety disorders (Miller et al., 1995), PTSD (King et al., 2013; Serpa et al., 2014), OCD (Didonna et al., 2019), stress, and general psychopathology (Alsubaie et al., 2017; Cásedas, 2021; Goleman & Davidson, 2018; Greeson et al., 2015; Riquelme-Marín et al., 2022; Sharpio & Weisbaum, 2020; Song & Lindquist, 2015).

Of the three fundamental skills cultivated through systematic meditation training (i.e., concentration, sensory clarity, and equanimity, Young, 2016) equanimity appears crucial to the health benefits of mindfulness meditation (Lindsay, Young, et al., 2018; Lord et al., 2025). Equanimity can be understood as the ability to maintain an open and accepting attitude towards all experience, whether pleasant, unpleasant, or neutral. It manifests as a combination of a dispositional trait or attitude, a dynamic phenomenological state, and a trainable cognitive skill (Desbordes et al., 2015; Lindsay & Creswell, 2017; Rogers et al., 2021). Mechanistically, equanimity is the system’s capacity to regulate the magnitude and persistence of affective, cognitive, and behavioral reactivity to primary sensory events and internal mental representations, independent of valence.

A dismantling trial conducted by Lindsay and colleagues (Lindsay, Chin, et al., 2018; Lindsay et al., 2019; Lindsay, Young, et al., 2018; Villalba et al., 2019) revealed that those engaging in equanimity supplemented with concentration, classified as the “Monitor + Accept” group, experienced greater reductions in stress, measured by cortisol, as well as greater improvements loneliness, social interaction, and general well-being compared to those engaging in concentration training alone. Given that acceptance or equanimity has been identified as a key cognitive factor on meditation’s wide-ranging impact on wellbeing, there is recent interest in exploring ways to enhance meditation by targeting equanimity-related brain processes with non-invasive neuromodulation (Lord et al., 2025).

In meditation, greater equanimity reduces default mode network-mediated self-referential processes (e.g., rumination, autobiographical elaboration, or self-evaluation), thus allowing experiences to arise and pass without amplification into prolonged distress or cognitive loops (Desbordes et al., 2015; Lord et al., 2025; Rahrig et al., 2022). Research has identified the posterior cingulate cortex (PCC), a primary midline hub of the default mode network (DMN; Andrews-Hanna et al., 2010), as a critical region for self-referential processing, especially the tendency to identify or “get caught up in” one’s experience (Brewer et al., 2013). Increased PCC activity has been associated with self-referential thought, while decreased activity has been associated with “effortless awareness” or being more present-minded (Brewer et al., 2013; Brewer & Garrison, 2014; Garrison et al., 2013). Increased activity within the PCC, along with heightened functional connectivity between the PCC and the subgenual cingulate cortex, has been shown to be potentially maladaptive, manifesting as rumination (e.g., repetitive negative thinking), which is a common symptom observed in clinical depression (Andrews-Hanna et al., 2014). Experienced meditators demonstrate an ability to willfully reduce activity in their PCC as they meditate (van Lutterveld et al., 2017). This control over their internal state is associated with increased anticorrelation between the DMN and the central executive network (CEN) (Bauer et al., 2019). These observations suggest that suppressive neuromodulation targeting the PCC could increase the dynamic flexibility of DMN and/or create more network segregation between DMN and CEN, thereby facilitating enhanced equanimity (Lord et al., 2025).

An ideal tool that holds promise for this approach is transcranial focused ultrasound (tFUS). tFUS can target subcortically (Kuhn et al., 2023), allowing for direct access to the PCC with optimal precision (Blackmore et al., 2019). tFUS has been shown to improve behavioral outcomes like reduction of depression and anxiety symptoms (Chou et al., 2024; Mahdavi et al., 2023). Previous research from Lord and colleagues showed that tFUS targeting the PCC reduced functional connectivity across the midline of the DMN, between the anterior medial prefrontal cortex and PCC, as well as increased state mindfulness and altered the sense of self and time (Lord et al., 2024).

As an extension of Lindsay and colleagues’ findings, (Lindsay, Chin, et al., 2018; Lindsay et al., 2019; Lindsay, Young, et al., 2018; Villalba et al., 2019), the present study aimed to assess the synergistic effects of combining meditation training targeting equanimity with tFUS on well-being. We hypothesized that suppressive (Dell’Italia et al., 2022) transcranial focused ultrasound (tFUS) targeting the posterior cingulate cortex (PCC) will produce an acute increase in equanimity, which, when combined with mindfulness training, will lead to greater improvements in well-being and measurable changes in functional connectivity in the brain. We used a single-blind, randomized design: participants received either all active or all sham tFUS stimulation across four sessions during their two weeks of meditation training. The effects were measured using self-report, mood scale, physiological, neuroimaging, and behavioral data.

## Methods

### Participants & Procedure

In total, 371 individuals expressed interest in the study through scanning a QR code on flyers in the Tucson area. Thirty-five individuals passed the initial phone screening. Exclusion criteria included experience with formal meditation training, mental health diagnosis, current user of psychotropic pharmaceuticals, history of epilepsy, heart disease, stroke, or migraines. Participants were English-speaking, right-handed, over 18 years old, and meditation naïve. Participants who passed these criteria were then scheduled for the first study visit. Seven eligible participants did not start the study, due to scheduling concerns. Twenty-four total participants completed the study, with 16 in the active tFUS group and 8 in the sham tFUS group. Out of the 24 participants who completed the study, 12 were female and 12 were male. The average age of participants was 26, with an age range of 18-53. The participant recruitment process is illustrated in the consort diagram in figure 1. Participants engaged in three weeks of study procedures, including six laboratory visits, two MRI sessions, 14 daily meditation lessons, and six days of ecological momentary assessments. The timeline is illustrated in figure 2.

**Figure 1.**
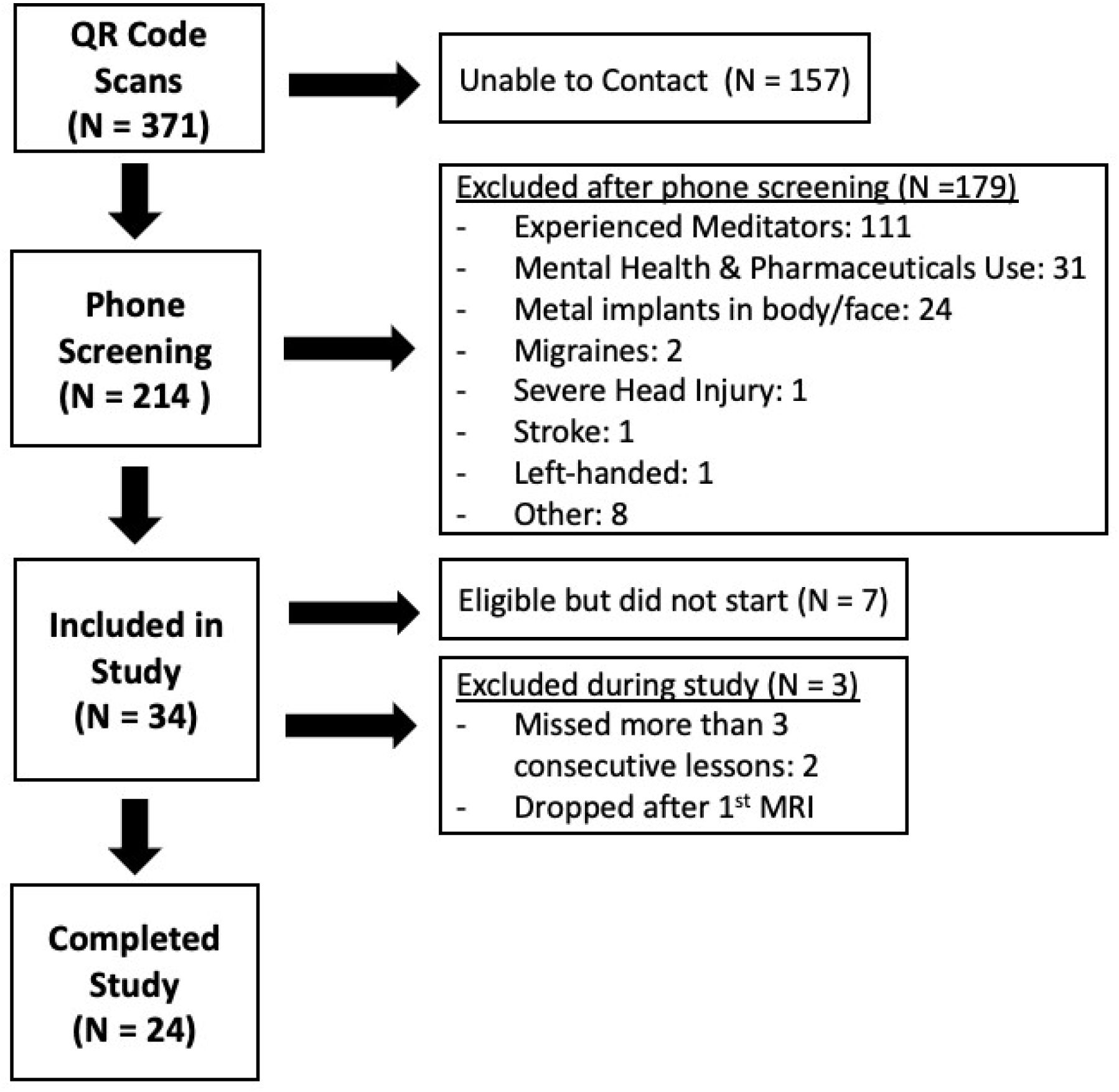
Consort diagram.

**Figure 2.**
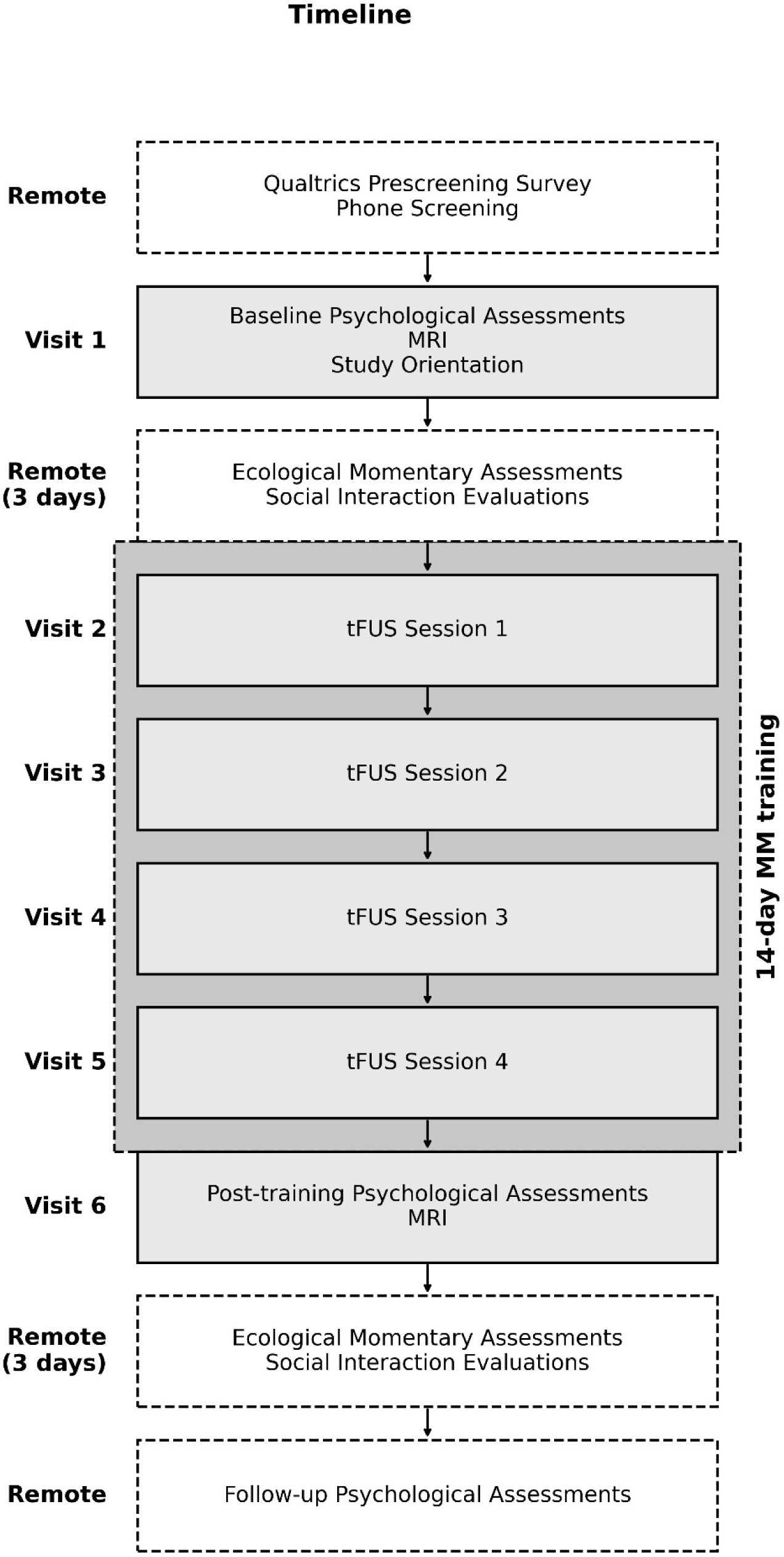
Study timeline. Participants performed baseline assessments before undergoing 14 consecutive days of daily at-home mindfulness meditation training, supplemented by in-person tFUS (active or sham) sessions. Post-intervention assessments were then made after the meditation training.

### MRI

#### Acquisition

A Siemens Skyra 3-Tesla scanner was used to acquire MR images. T1-weighted anatomical images (MP-RAGE; TR=2300 ms; TE=2.32 ms; TI=900 ms; flip angle=8; FOV=240 mm x 240 mm) and functional BOLD scans were acquired (TR=2000 ms; TE=30 ms; flip angle=90; FOV=1920 mm x 1920 mm; voxel size 1.5 mm x 1.5 mm x 1.5 mm). Susceptibility-Weighted Imaging (SWI) scans were also acquired (TR=28 ms; TE=20 ms; flip angle=15; FOV=220 mm x 178.75 mm). During the resting-state BOLD scan, participants were asked to look at a fixation cross. After the meditation training was completed, another sequence of MR images was acquired. Two BOLD scans were acquired with the same parameters used previously. In the first scan, participants were asked to stare at a fixation cross in the same manner. In the second scan, participants were asked to perform the body focus meditation they learned in their lessons. A second SWI scan was also acquired to assess if any damage or changes to vasculature occurred during the training period. MR data were preprocessed using a standard CONN pipeline (Whitfield-Gabrieli & Nieto-Castanon, 2012) (RRID:SCR_009550) release 22.a (Nieto-Castanon & Whitfield-Gabrieli, 2021). Details are provided in Supplemental Materials.

#### First-level analysis

Regions of interest (ROIs) corresponding to the DMN (21 ROIs across 3 subnetworks) and CEN (16 ROIs across 3 subnetworks) were identified using a 17-network, 100-parcel atlas (Schaefer et al., 2018), in order to capture large-scale interactions across functional networks. ROI-to-ROI connectivity values were computed using the CONN toolbox’s ROI-to-ROI pipeline, which applies Fisher-transformed bivariate correlation coefficients to estimate functional coupling between regions. For each participant, pairwise connectivity estimates were extracted between all DMN and all CEN ROIs. To assess overall DMN-CEN connectivity strength, these values were averaged to yield a single summary index for each scan, yielding baseline and post-intervention values for each participant. To assess subnetwork connectivity strength, the relevant ROIs were averaged together.

### tFUS Sessions

#### Targeting

A custom-written MATLAB script was used to determine the optimal functional imaging-based target location for the tFUS, which was defined as the peak DMN activity in the PCC. To determine this, ICA was performed on preprocessed BOLD data using the Group ICA/IVA Toolbox (GIFT; version 1.3.8; Calhoun et al., 2001). To identify the DMN components, spatial regression was performed using a DMN template (Hu et al., 2016). Because the DMN was sometimes distributed across multiple components, the final DMN component was obtained by summing the normalized weight of the regression values of the components. This DMN component was then masked to only the PCC region using a template (Hu et al., 2016; Yang et al., 2014), and the peak value voxel was located. These coordinates were converted to MNI coordinates for translation to the structural image. When the structural data was being loaded into the neuronavigation software (visor2), Talairach coordinates were identified so that these target MNI coordinates could be transformed to the raw structural image for accurate neuronavigation.

#### tFUS Administration

Participants were randomized to either the stimulation group or the sham group. The tFUS parameters are outlined below in table 1. A function generator (BK Precision 4078 waveform generator) controlled by a computer script created the ultrasound signal that was driven through an amplifier (E&I 210L power amplifier) to a single-element fixed-depth 55 mm focused ultrasound transducer (Blatek AT35246), coupled to the scalp using ultrasound transmission gel. Based on Dell’Italia and colleagues (2022), a 5% duty cycle was selected for its suppressive effects. See table 1 for full tFUS parameters. Once the participant’s head was registered to the neuronavigation software (ANT Neuro visor2), they were asked to close their eyes and begin meditating while a study technician held the transducer in place by hand for the 5-minute duration of the ultrasound delivery. For participants receiving sham stimulation, the function generator’s oscillators were turned off so that no signal was produced or emitted. After receiving tFUS, participants were escorted to a meditation room and were instructed to meditate for as long as they would like. The meditation duration was recorded after each visit. When participants returned from mediating, they completed a sensation questionnaire to facilitate a reporting system in case of any adverse events.

**Table 1:** tFUS Parameters.

**tFUS Parameters**
| **Parameter** | **Value** |
| --- | --- |
| Acoustic frequency | 500 kHz |
| Duty Cycle | 5% |
| Stimulus Pattern | 5 seconds on, 10 seconds off for 5 minutes |
| Target | Peak PCC activity in DMN |
| I<sub>SPTA</sub> | 1938.61 (mW/cm<sup>2</sup>) |
| I<sub>SPPA</sub> | 3.88 (W/cm<sup>2</sup>) |

### Mindfulness Meditation Training Protocol

The mindfulness meditation training protocol was the same 14-day Body Focus training that was used in the Lindsay et. al., studies. A link to the lesson was sent to the participant’s cell phone via text message using Qualtrics each day. The participants completed the first day of the program in a meditation room set up in the Science Enhanced Mindful Awareness (SEMA) Lab at the University of Arizona to ensure the participant’s initial understanding of the training before completing the rest of the sessions on their own. The training material was released to participants every 24 hours via text message, and they had the ability to rewatch lessons if needed. Participants were not allowed to move onto the next lesson until they finished the current session, nor could they perform future days’ lessons ahead of schedule. If a participant missed a meditation lesson, they were sent a reminder. Participants were allowed to perform multiple lessons a day to catch up. However, if they missed more than three sessions, they were dropped from the study.

## Results

### Adherence

Three participants withdrew during the study; two of these were due to not being able to keep up with daily lessons, and one dropped out before the first lesson due to scheduling constraints. No participants withdrew due to adverse effects. Twenty-four people completed the study.

### tFUS Meditation Sessions

No adverse events were reported. In 40 out of 56 active sessions recorded (13 out of 14 participants; the first 2 participants did not complete sensation surveys), participants reported hearing the tFUS. This was usually described as a “beeping” or a “buzzing” or a “high-pitched tone,” which likely corresponds with the audible 1000 Hz PRF. In 5 out of 32 sham sessions (3/8 participants), the participant reported hearing the tFUS. These sounds were described as a “whirring” or a “start up” noise, which likely corresponds to the sound of the amplifier being turned on just prior to tFUS administration. Data from the sensation survey are presented in figure 3. Most sessions yielded few to zero sensations reported.

**Figure 3.**
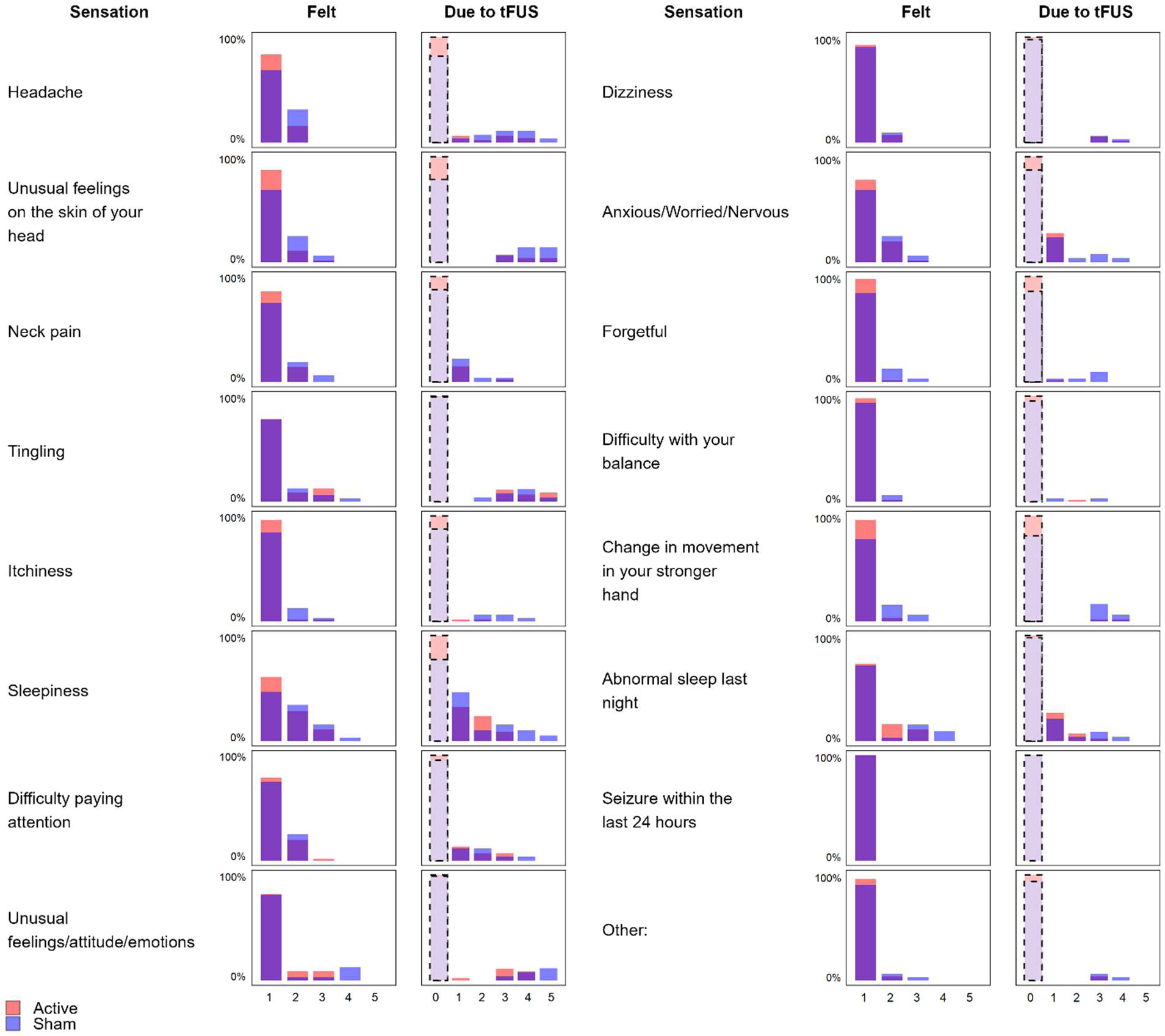
Survey of sensations felt after tFUS meditation sessions. Histograms for each row show, on the left, percentages for ratings of how much each sensation was felt on a scale of 1-5, and on the right, how much this sensation is attributed to the tFUS. Lighter, dash-outlined bars in the “due to tFUS” column correspond to instances in which the sensation was rated as being not present, meaning that there is nothing to attribute to tFUS in the first place. Red marks active condition, and blue marks sham. The color appears purple when they overlap. The first 2 participants did not complete the sensation survey.

### MRI

#### DMN-CEN Connectivity Change

A linear mixed-effects model was fit to examine the effects of Condition (Active vs. Sham), Session (Pre vs. Post), and their interaction on DMN-CEN functional connectivity, with a random intercept for participant to account for repeated measures (48 observations from 24 participants). The analysis revealed a robust Condition × Session interaction (β = −0.118, SE = 0.033, t(44) = −3.61, p < 0.001, 95% CI [−0.184, −0.052]), indicating that connectivity changed in opposite directions across the session in the two conditions (see figure 4). DMN-CEN connectivity reduced significantly in the Active condition (slope = −0.067, 95% CI [−0.104, - 0.030], p < 0.001), while it increased in the Sham condition (slope = +0.051, 95% CI [−0.001, 0.103], p = 0.055). There was no significant difference between the two conditions at baseline. See table 2.

**Figure 4.**
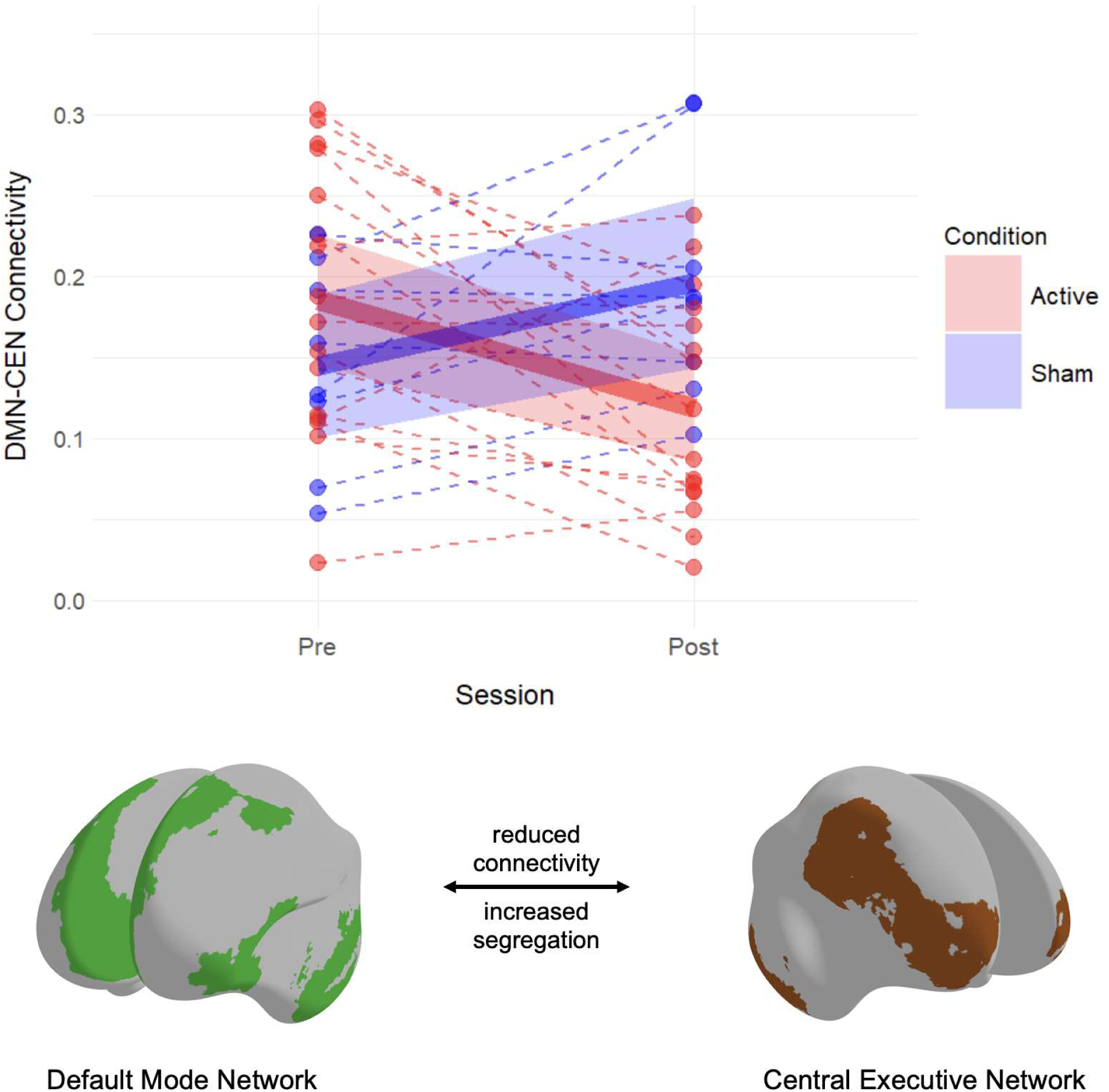
Functional connectivity between the DMN and CEN as a function of condition and session. Individual participants are shown with dots and dashed lines (Active = red; Sham = blue). Bold lines represent estimated marginal means with 95% confidence bands. A significant Condition × Session interaction was observed (p < 0.001), reflecting a pre-to-post decrease in connectivity in the Active condition (slope = −0.067, 95% CI [−0.104, −0.030]) and a slight increase in Sham (slope = +0.051, 95% CI [−0.001, 0.103]).

**Table 2:** Fixed effects from the mixed model for DMN-CEN functional connectivity.

| Effect | Estimate | SE | *p*-value | 95% CI |
| --- | --- | --- | --- | --- |
| Intercept | 0.145 | 0.025 | <0.001** | [0.094, 0.195] |
| Condition | 0.041 | 0.031 | 0.185 | [-0.021, 0.103] |
| Session | 0.051 | 0.027 | 0.055 | [-0.001, 0.103] |
| Condition × Session | -0.118 | 0.033 | <0.001** | [-0.184, -0.052] |

*Note.* \*\* $p < 0.001$ ***DMN-CEN Subnetwork Connectivity Changes***

#### DMN-CEN Subnetwork Connectivity Changes

For each DMN-CEN edge within the three DMN and three CEN subnetworks, a linear mixed-effects model with fixed effects for Condition and Session and a participant random intercept was fit, and p-values were adjusted for multiple comparisons using the Benjamini–Hochberg false-discovery procedure. All edges showed a negative estimate, suggesting a broad segregation, and significant effects were observed in all edges involving either the DMN_A_ or CEN_B_. See figure 5 and table 3.

**Figure 5.**
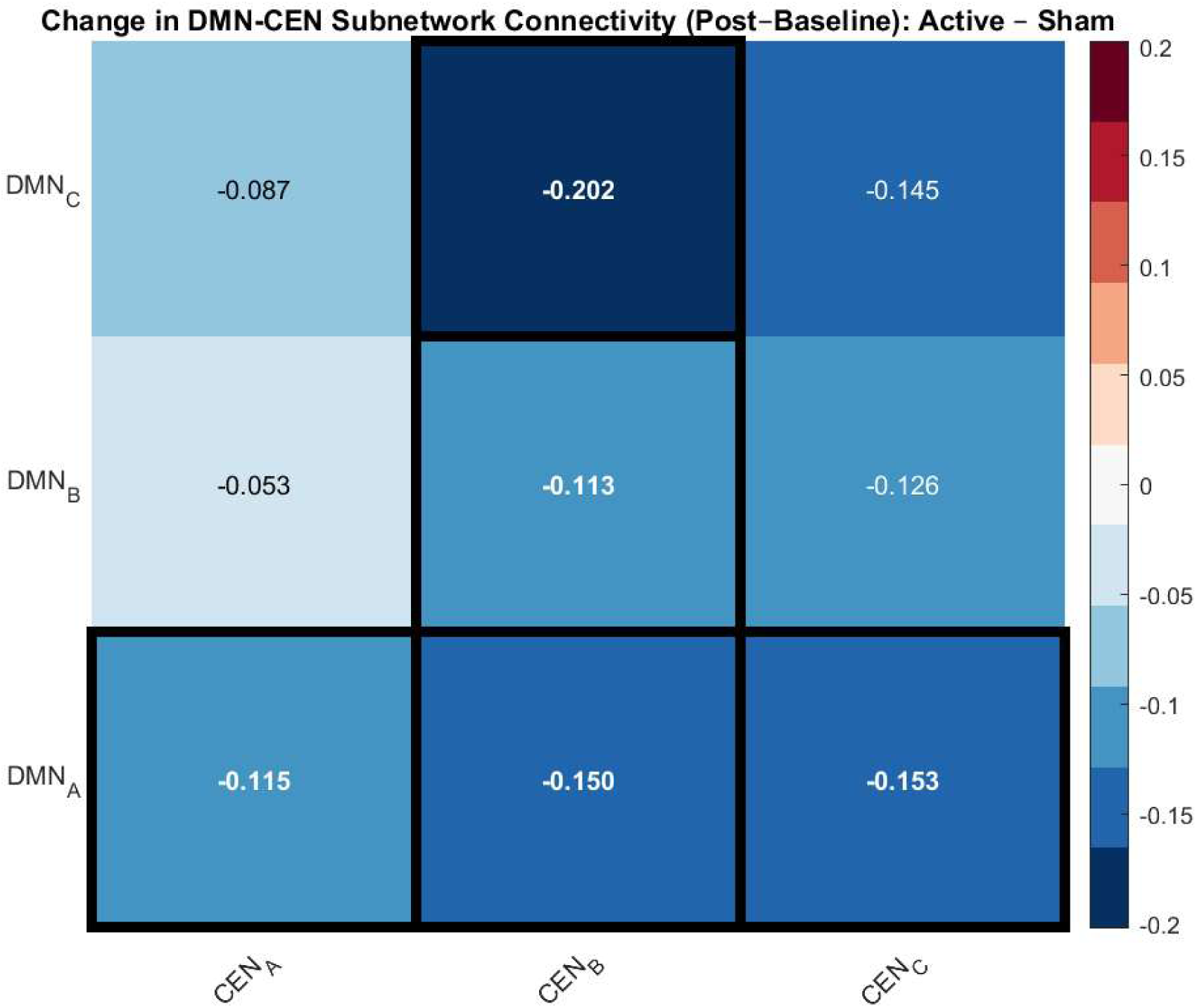
Group differences in post–baseline changes in functional connectivity between DMN and CEN subnetworks. Each cell displays the mean Fisher-z–transformed interaction effect of *Condition × Session* (Active − Sham) from the linear mixed-effects model estimated for all ROI-to-ROI pairs between a given DMN (rows) and CEN (columns) subnetwork. Negative values indicate greater DMN–CEN decoupling in the Active condition following the intervention. Black outlines mark subnetworks showing significant group differences after correcting for multiple comparisons using the Benjamini-Hochberg method.

**Figure 6.**
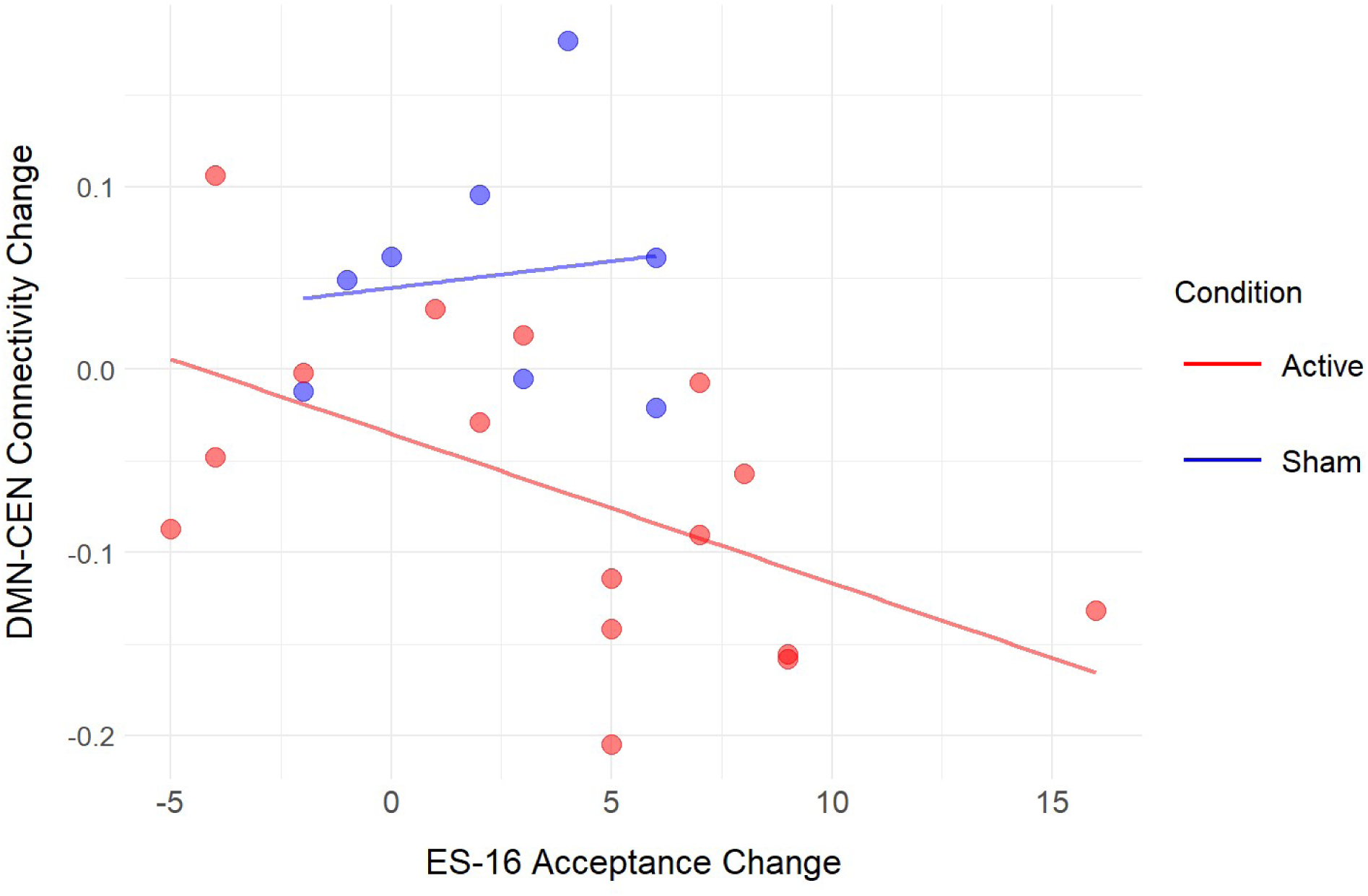
Relationship between change in ES-16 Acceptance and change in resting state DMN-CEN functional connectivity. Each point reflects one participant (Active = red; Sham = blue). Regression lines depict condition-specific slopes. In the Active condition, greater increases in Acceptance tended to correspond with greater decreases in DMN-CEN connectivity, whereas in Sham there was no relationship. The slope difference (interaction) was large but not significant (Δβ = 44.1, p = 0.11).

**Figure 7.**
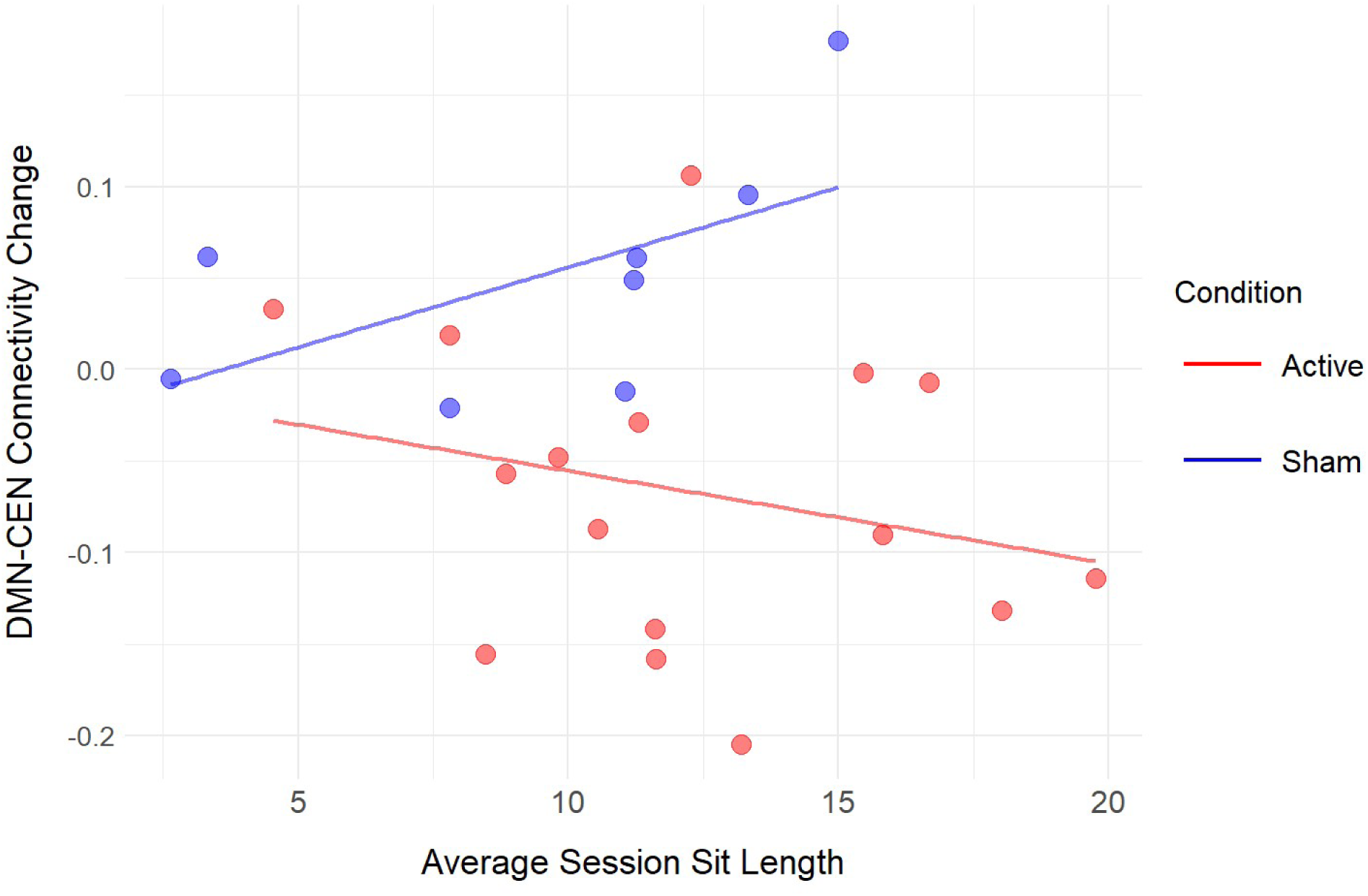
Relationship between meditation practice (average session sit length, in minutes) and DMN-CEN connectivity change. Each point reflects one participant (Active = red; Sham = blue). Regression lines depict condition-specific slopes. In the Active condition, longer average meditation sessions tended to correspond with greater decreases in DMN-CEN connectivity, whereas in Sham the slope trended in the opposite direction. The slope difference (interaction) was statistically significant (p = 0.042).

**Table 3.** DMN-CEN subnetwork changes.

| **DMN**<br>**Subnetwork** | **CEN**<br>**Subnetwork** | **Model**<br>**Coefficient** | ***p*-uncorrected** | ***p*-FDR**<br>**corrected** |
| --- | --- | --- | --- | --- |
| DMN<sub>A</sub> | CEN<sub>A</sub> | -0.115 | 0.024 | 0.004* |
| DMN<sub>A</sub> | CEN<sub>B</sub> | -0.150 | 0.009 | 0.028* |
| DMN<sub>A</sub> | CEN<sub>C</sub> | -0.153 | 0.005 | 0.022* |
| DMN<sub>B</sub> | CEN<sub>A</sub> | -0.054 | 0.349 | 0.349 |
| DMN<sub>B</sub> | CEN<sub>B</sub> | -0.113 | 0.020 | 0.043* |
| DMN<sub>B</sub> | CEN<sub>C</sub> | -0.126 | 0.117 | 0.150 |
| DMN<sub>C</sub> | CEN<sub>A</sub> | -0.087 | 0.190 | 0.213 |
| DMN<sub>C</sub> | CEN<sub>B</sub> | -0.202 | 0.002 | 0.018* |
| DMN<sub>C</sub> | CEN<sub>C</sub> | -0.145 | 0.043 | 0.065 |

*Note.* \* $p$ -FDR<0.05.

A more granular analysis of the individual ROIs was performed using the same method. No edges reached the threshold of significance, once corrections for multiple comparisons were made. See figure S1 in supplementary material.

#### Brain-behavior link with acceptance

A linear model relating changes in Acceptance to changes in DMN-CEN connectivity revealed a significant overall negative association (β = −22.9, SE = 9.5, t(22) = −2.41, p = 0.025), such that greater decreases in connectivity predicted greater increases in Acceptance. When Condition was included, the association was significant in the Active group (β = −38.0, SE = 13.8, p = 0.012) but not in the Sham group (β = +6.1, SE = 19.6, p = 0.76). The slope difference between conditions was large but nonsignificant (Δβ = 44.1, p = 0.11).

#### Brian-behavior link with meditation sitting time

To test whether active tFUS impacted the relationship between meditation practice and neural change, a linear model was fit with average meditation session length (minutes across four sessions) as the outcome and DMN-CEN connectivity change (Post - Pre), Condition (Active vs. Sham), and their interaction as predictors (N = 24; 16 Active, 8 Sham). The model indicated a significant DMN-CEN × Condition interaction (β = −51.69, SE = 23.79, t(20) = −2.17, p = 0.042), demonstrating that the association between connectivity change and practice length differed reliably by condition.

In the Sham group, longer average session lengths corresponded with a nonsignificant trend of increases in DMN-CEN connectivity (β = +39.72, SE = 20.90, p = 0.072, 95% CI [−3.88, 83.31]), while in the Active group, longer practice was associated with decreases in connectivity (β = −11.98, SE = 11.37, p = 0.305, 95% CI [−35.69, 11.73]). Although neither slope was significant on its own, the significant interaction confirmed a divergence between conditions.

## Discussion

The present study investigated the use of tFUS to augment a two-week at-home meditation training course, with the goal of amplifying the effects of mindfulness meditation using an individualized tFUS-targeting approach. There were significant changes in resting state BOLD functional connectivity as a result of this intervention. Active tFUS produced robust decreases in resting state DMN-CEN connectivity relative to sham, with connectivity decoupling in the stimulation group but rising slightly in controls. A closer analysis revealed that while these changes were broadly exhibited across all subnetworks and most individual ROIs, they were statistically significant when edges connected with either the subnetworks DMN_A_ or CEN_B_. Notably, these large-scale network effects were behaviorally meaningful, as greater decreases in DMN-CEN connectivity corresponded with both larger gains in acceptance as measured by the ES-16 and longer time sitting in tFUS meditation sessions.

Increased anticorrelation between DMN and CEN is a hallmark of successful meditation practice and the acquisition of trait-level shifts in mindfulness (Bauer et al., 2019; Cásedas, 2021; Goleman & Davidson, 2018; Treves et al., 2024), typically interpreted as enhanced top-down regulation of internal mentation and mind-wandering. These network connectivity shifts are observed over hundreds of hours or more of reported meditation practice, and we observed a shift in this direction in just two weeks. Moreover, the same pattern did not merely emerge more quickly: the Condition x Session effect showed that DMN-CEN coupling decreased in the active group but increased in sham, indicating that the active tFUS redirected the training trajectory rather than merely accelerating it, and it redirected it in the direction of what is seen after hundreds of hours of practice.

The subnetwork effects clarify the mechanisms at play. The DMN_A_ contains the core (PCC/precuneus ↔ anterior PFC) of the DMN, and activity here is associated with self-referential thought and internal mentation (Andrews-Hanna, 2012; Andrews-Hanna et al., 2010; Buckner & DiNicola, 2019; Schaefer et al., 2018). The CEN_B_ (bilateral lPFC ↔ right IPL and lateral temporal) corresponds to the externally-oriented control system that preferentially couples with the dorsal attention network (DAN) and salience network (SN) for attention and task control (Dixon et al., 2018). The DMN_A_ and CEN_B_ decoupling shows that the active tFUS thus promoted a decentering of self-referential cores from externally directed control systems, consistent with a shift toward present-oriented, stimulus-guided attention that is mechanistically distinct from practice-alone effects.

This mechanistic specificity is important: if the active tFUS simply enhanced or accelerated training, both groups would have changed in the same direction with larger magnitude in active. Instead, the change in DMN-CEN connectivity diverged between the two conditions, indicating a qualitatively different network reconfiguration induced by tFUS plus meditation training. This can be made sense of in the context that the acquisition of cognitive skills generally follows an inverted u-shaped curve in the associated brain activity, and systematic focus training (meditation) is no different (Brefczynski-Lewis et al., 2007). Novices often show a muddled segregation between DMN and CEN (Bauer et al., 2019; Josipovic et al., 2012); early attempts at meditation may even show a slight increase in self-referential activity as they get stuck in the mind-wandering phase (DMN) of their fluctuating cognitive process (Brewer et al., 2011; Hasenkamp et al., 2012). Therefore, aiding individuals through the initial difficulty of quieting restless mind-wandering activity may greatly accelerate their long-term acquisition of these skills.

Despite the considerable burden on participants (six total lab visits, numerous surveys, and daily training), they showed high compliance. This may have partly been achieved by asking them at the beginning of the study why they were interested in learning meditation and what they hoped to get out of the lessons they were signing up for. This was intentionally designed to create a meaningful “buy-in” from participants, so they would have self-generated reasons to push through the course.

### Limitations & Future Directions

There were several limitations in this study. The sample size was much lower (n=24 vs. n=144) than the meditation training study that was used as its basis, which greatly constrains its statistical power and generalizability (Lindsay, Young, et al., 2018). Although an individualized protocol was used to target the PCC peak within the DMN, no correction for skull aberration was applied. Consequently, some participants may have received suboptimal energy at the PCC target due to the skull distorting and attenuating the beam, although the effect of the skull would be to shorten the focal length, thus targeting the PCC-precuneus junction (and some research supports the precuneus as part of the DMN). Future individualized applications of tFUS should incorporate acoustic modeling (e.g., CT- or pseudo-CT simulations) and, if possible, distortion correction to maximize the accuracy of the tFUS exposure. Integrating these steps into the neuronavigation workflow would extend the structural and functional based targeting used here to a higher level of precision.

Survey instruments for mindfulness and equanimity also warrant caution. As novices practice, response shifts can occur. Increased internal clarity may lead participants to endorse more discomfort or lower equanimity despite genuine skill growth. Future assessments can improve on this by employing targeted phenomenological interviews to capture greater depth and detail into the practitioner’s experience, allowing important experiences to be correctly contextualized.

It may also be the case that the approach of targeting the PCC with suppressive tFUS is not appropriate or optimal for every individual seeking to accelerate the acquisition of systematic focus training skills. Future research should endeavor to characterize what a given individual needs most, from both a neurobiological and behavioral perspective, including the optimal dose or exposure time to this type of tFUS. We have previously advocated for this “precision wellness” approach as a promising direction for these types of interventions (Lord et al., 2025).

## Supplemental Materials

### MRI Preprocessing

#### Preprocessing

Results included in this manuscript come from analyses performed using CONN (Whitfield-Gabrieli & Nieto-Castanon, 2012) (RRID:SCR_009550) release 22.a (Nieto-Castanon & Whitfield-Gabrieli, 2021) and SPM (Penny et al., 2011) (RRID:SCR_007037) release 12.7771.

Functional and anatomical data were preprocessed using a flexible preprocessing pipeline (Nieto-Castanon, 2020b) including realignment with correction of susceptibility distortion interactions, slice timing correction, outlier detection, direct segmentation and MNI-space normalization, and smoothing. Functional data were realigned using SPM realign & unwarp procedure (Andersson et al., 2001), where all scans were coregistered to a reference image (first scan of the first session) using a least squares approach and a 6 parameter (rigid body) transformation (Friston et al., 1995), and resampled using b-spline interpolation to correct for motion and magnetic susceptibility interactions. Temporal misalignment between different slices of the functional data (acquired in interleaved Siemens order) was corrected following SPM slice-timing correction (STC) procedure (Henson et al., 1999; Sladky et al., 2011), using sinc temporal interpolation to resample each slice BOLD timeseries to a common mid-acquisition time. Potential outlier scans were identified using ART (Whitfield-Gabrieli et al., 2011) as acquisitions with framewise displacement above 0.9 mm or global BOLD signal changes above 5 standard deviations (Nieto-Castanon, submitted; Power et al., 2014), and a reference BOLD image was computed for each subject by averaging all scans excluding outliers. Functional and anatomical data were normalized into standard MNI space, segmented into grey matter, white matter, and CSF tissue classes, and resampled to 2 mm isotropic voxels following a direct normalization procedure (Calhoun et al., 2017; Nieto-Castanon, submitted) using SPM unified segmentation and normalization algorithm (Ashburner, 2007; Ashburner & Friston, 2005) with the default IXI-549 tissue probability map template. Last, functional data were smoothed using spatial convolution with a Gaussian kernel of 8 mm full width half maximum (FWHM).

#### Denoising

In addition, functional data were denoised using a standard denoising pipeline (Nieto-Castanon, 2020a) including the regression of potential confounding effects characterized by white matter timeseries (5 CompCor noise components), CSF timeseries (5 CompCor noise components), motion parameters and their first order derivatives (12 factors) (Friston et al., 1996), outlier scans (below 26 factors) (Power et al., 2014), session and task effects and their first order derivatives (6 factors), and linear trends (2 factors) within each functional run, followed by bandpass frequency filtering of the BOLD timeseries (Hallquist et al., 2013) between 0.008 Hz and 0.09 Hz. CompCor (Behzadi et al., 2007; Chai et al., 2012) noise components within white matter and CSF were estimated by computing the average BOLD signal as well as the largest principal components orthogonal to the BOLD average, motion parameters, and outlier scans within each subject’s eroded segmentation masks. From the number of noise terms included in this denoising strategy, the effective degrees of freedom of the BOLD signal after denoising were estimated to range from 103.3 to 118.1 (average 113.4) across all subjects (Nieto-Castanon, submitted).

---

**Table S1.**
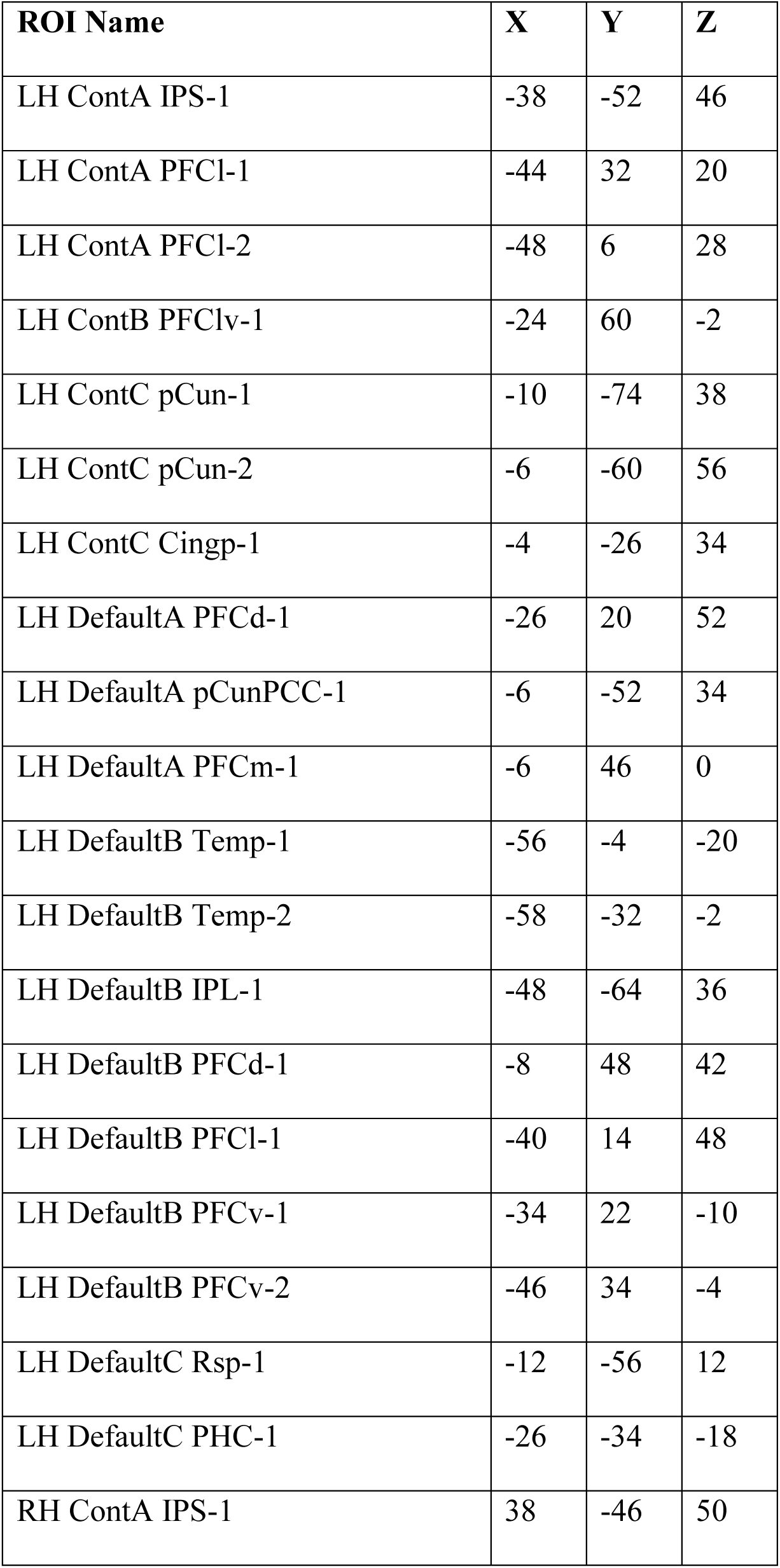
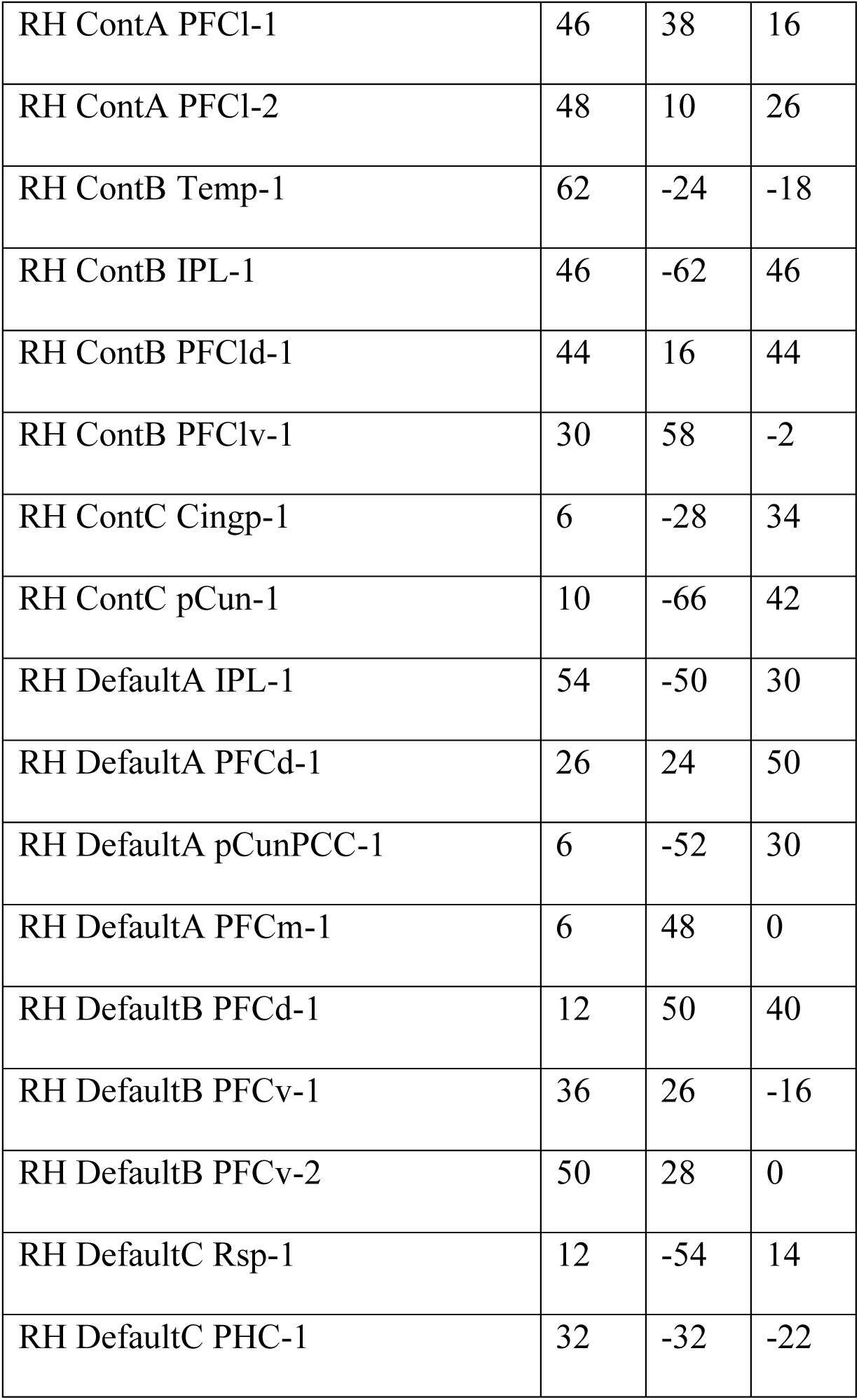
ROI MNI Centroids.

**Figure S1.**
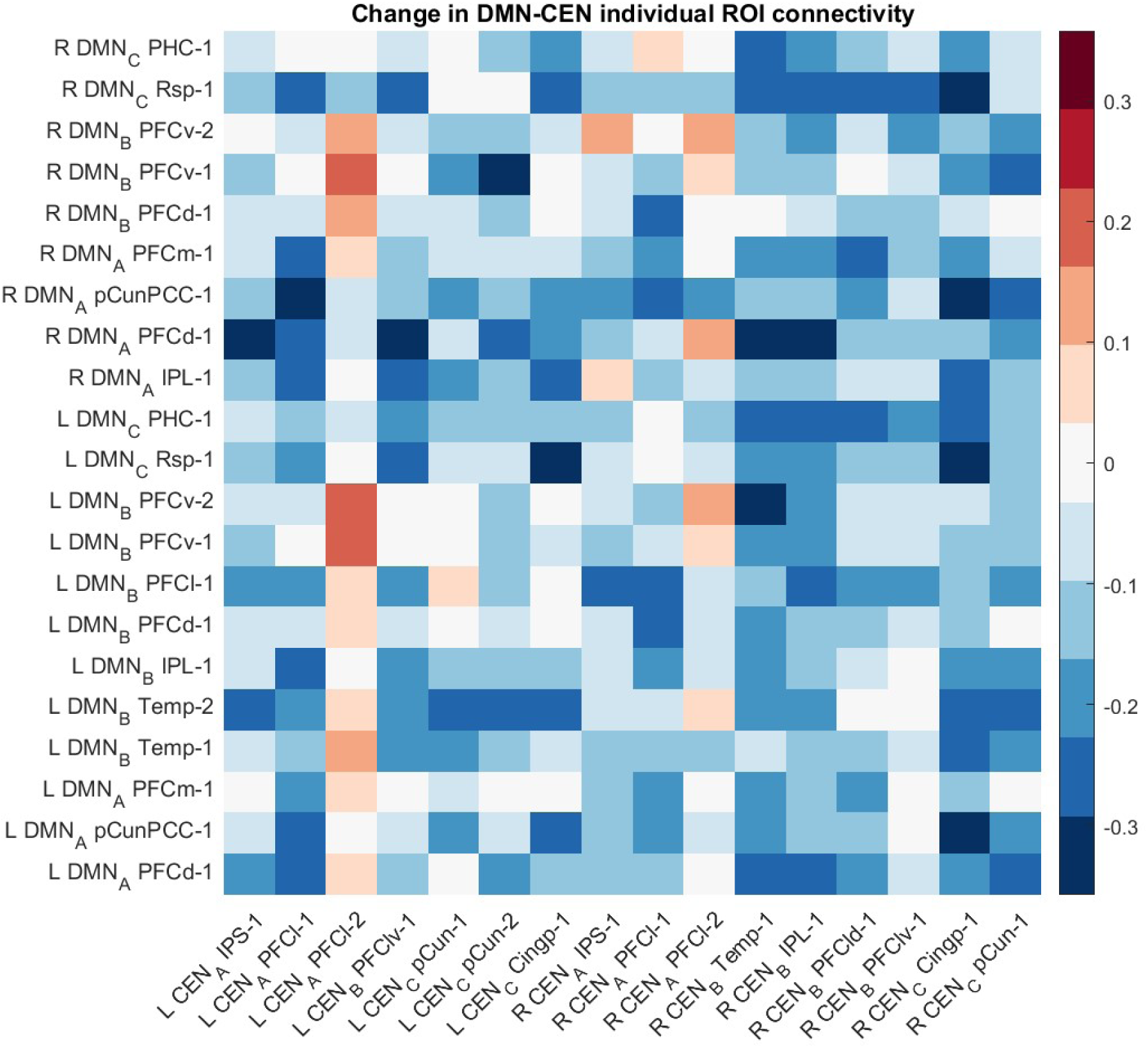
Group differences in post–baseline changes in functional connectivity between DMN and CEN individual ROIs. Each cell displays the mean Fisher-z–transformed interaction effect of *Condition × Session* (Active − Sham) from the linear mixed-effects model estimated for all ROI-to-ROI pairs between a given DMN (rows) and CEN (columns) ROI. Negative values (cooler colors) indicate greater DMN–CEN decoupling in the Active condition following the intervention, while positive values (warmer colors) indicate greater coupling.

## Notes

### Competing Interest Statement

Author Joseph L Sanguinetti receives salary and is a shareholder in Sanmai Technologies. Author Shinzen Young receives consulting fees from Sanmai Technologies. No other competing interests.

### Funding

- Joy Ventures

